# An Iron-regulated Signalling Pathway Controls Adipose Browning and Cancer Cachexia

**DOI:** 10.1101/2025.07.16.664180

**Authors:** Jung Seung Nam, Maya S. Dixon, Pardis Ahmadi, Kathrin Schilling, Samuel Pan, Yuxuan Chen, Nal Ae Yoon, Yanping Sun, Jordan Lu, Shu Ichimiya, Jeanine M. Genkinger, Thomas C Caffrey, Kelsey A. Klute, Benjamin J. Swanson, Paul Grandgenett, Michael A. Hollingsworth, Kazuki Sugahara, Michael D. Kluger, Anthony Ferrante, Sabrina Diano, Iok In Christine Chio

## Abstract

The browning and atrophy of white adipose tissue (WAT) are early events in cachexia, a lethal metabolic disorder affecting nearly half of cancer patients, including those with pancreatic ductal adenocarcinoma (PDA). Using patient-derived specimens and PDA mouse models, we identified perturbations in iron metabolism and proteinaceous methionine oxidation as key initiating events of adipose browning. In particular, the iron influxes that accompany WAT browning induce the activity of methionine sulfoxide reductase A (MSRA), an enzyme that reverses the oxidation of proteinaceous methionine residues. Mechanistically, iron coordination by the conserved iron-binding motifs (E203-xx-H206) of two MSRA polypeptides serves to multimerize, stabilize, and enzymatically activate MSRA. This in turns facilitates adipose browning by maintaining the reduced state of two methionines near the ATP-binding site of Protein Kinase A (PKA). Remarkably, in mouse models of PDA, *MsrA* deletion impairs WAT browning, significantly mitigates cachexia, and improves the overall survival of tumor-bearing animals. By establishing the iron-MSRA-PKA axis as a key nexus of cancer-associated cachexia, our study offers new perspectives for the treatment of this condition.

## INTRODUCTION

Cancer patients who develop cachexia, a metabolic disorder characterized by morbid wasting of adipose tissue and skeletal muscle, suffer involuntary weight loss, poor responses to therapy, and elevated mortality rates (Baracos et al., 2018). While cachexia affects nearly half of cancer patients, it is especially prevalent (∼ 85% of cases) among those with pancreatic ductal adenocarcinoma (PDA), the most lethal pancreatic malignancy (Poulia et al., 2020). Although the biological factors that induce cachexia remain poorly defined, recent studies indicate that adipose tissue wasting precedes muscle loss in cancer patients (Agustsson et al., 2007; Sah et al., 2019; Zuijdgeest-van Leeuwen et al., 2000) and that muscle mass can be preserved in tumor-bearing animals by experimental blockage of adipose depletion (Das et al., 2011; Kir et al., 2014). Thus, elucidating the factors that initiate adipose loss should reveal the underlying mechanisms of cancer cachexia and provide new clinical strategies to forestall its development.

Mammals possess both white adipose tissue (WAT), which stores energy in the form of triacylglycerides, and brown adipose tissue (BAT), which can release energy in the form of heat. WAT, the predominant form of fat tissue in adults, is comprised mostly of white adipocytes that have a unilocular morphology marked by a single massive lipid droplet. With relatively few mitochondria, these cells have limited capacity for oxidative phosphorylation. In contrast, the brown adipocytes of BAT, which possess multiple small (multilocular) lipid droplets and abundant mitochondria, can robustly support oxidative phosphorylation. Also, brown adipocytes uniquely express high levels of uncoupling protein 1 (UCP1), which allows the proton motive force of mitochondria to be dissipated as heat (rather than ATP synthesis) in a physiological process known as non-shivering thermogenesis (Cannon and Nedergaard, 2004).

Under certain physiological conditions, such as cold exposure, WAT cells can be transformed into “beige adipocytes” that have BAT-like features, including a multilocular morphology, high mitochondrial content, and inducible UCP1 expression (Nedergaard and Cannon, 2014). As a result of this “browning” process, beige adipocytes gain a thermogenic potential that is readily induced by sympathetic tone. In particular, norepinephrine released by parenchymal neurites can activate β-adrenergic signaling in beige adipocytes to mobilize their lipid stores for thermogenesis (Cannon and Nedergaard, 2004; Cero et al., 2021; Hsieh and Carlson, 1957). Importantly, recent work has established that extensive WAT browning is a characteristic feature of cancer cachexia that antecedes muscle wasting (Kir et al., 2014; Neves et al., 2016; Petruzzelli et al., 2014; Zimmers et al., 2016). Indeed, adipose depletion, coinciding with rising body temperature, can already be observed in PDA patients months prior to diagnosis (Sah et al., 2019), consistent with its postulated role as an early trigger of cancer cachexia.

## RESULTS

### Methionine oxidation is altered during WAT browning in PDA cachexia

Previous studies have shown that PDA-bearing mice develop a cachectic phenotype marked by profound loss of muscle and adipose tissue (Michaelis et al., 2017; Talbert et al., 2019). Thus, to identify factors that induce adipose browning during cancer cachexia, we employed the autochthonous KPC (*Kras^G12D^; p53^R172H^; PdxCre*)(Hingorani et al., 2005) mouse model of PDA. As expected (Michaelis et al., 2017; Talbert et al., 2019), PDA-bearing KPC mice (mPDA) lost significant weight, with depletion of both adipose and lean mass, relative to age-matched non-tumor (mNT) controls (Fig. 1a). Adipose atrophy occurred to the same degree in male and female mice upon adjusting for body mass differences (Fig. S1a). The WAT adipocytes of KPC mice are markedly shrunken in size and exhibit a BAT-like multilocular morphology (Fig. 1b). These changes are accompanied by increased expression of *Ucp1* mRNA (Fig. S1b) and protein (Fig. S1c) in both epididymal and subcutaneous WAT (eWAT and sWAT). In normal mice, UCP1-mediated energy expenditure occurs in response to specific physiological stimuli, such as cold exposure (Nedergaard and Cannon, 2014). However, when monitoring mice maintained at ambient temperature, we also observed increased energy expenditures in tumor-bearing animals relative to age-matched controls. These increases reached statistical significance during the sleep cycle (Fig. 1c), even though activity levels (Fig. S1d), food intake (Fig. S1e), and water intake (Fig. S1f) were not significantly different between tumor-bearing and control mice.

**Figure 1.**
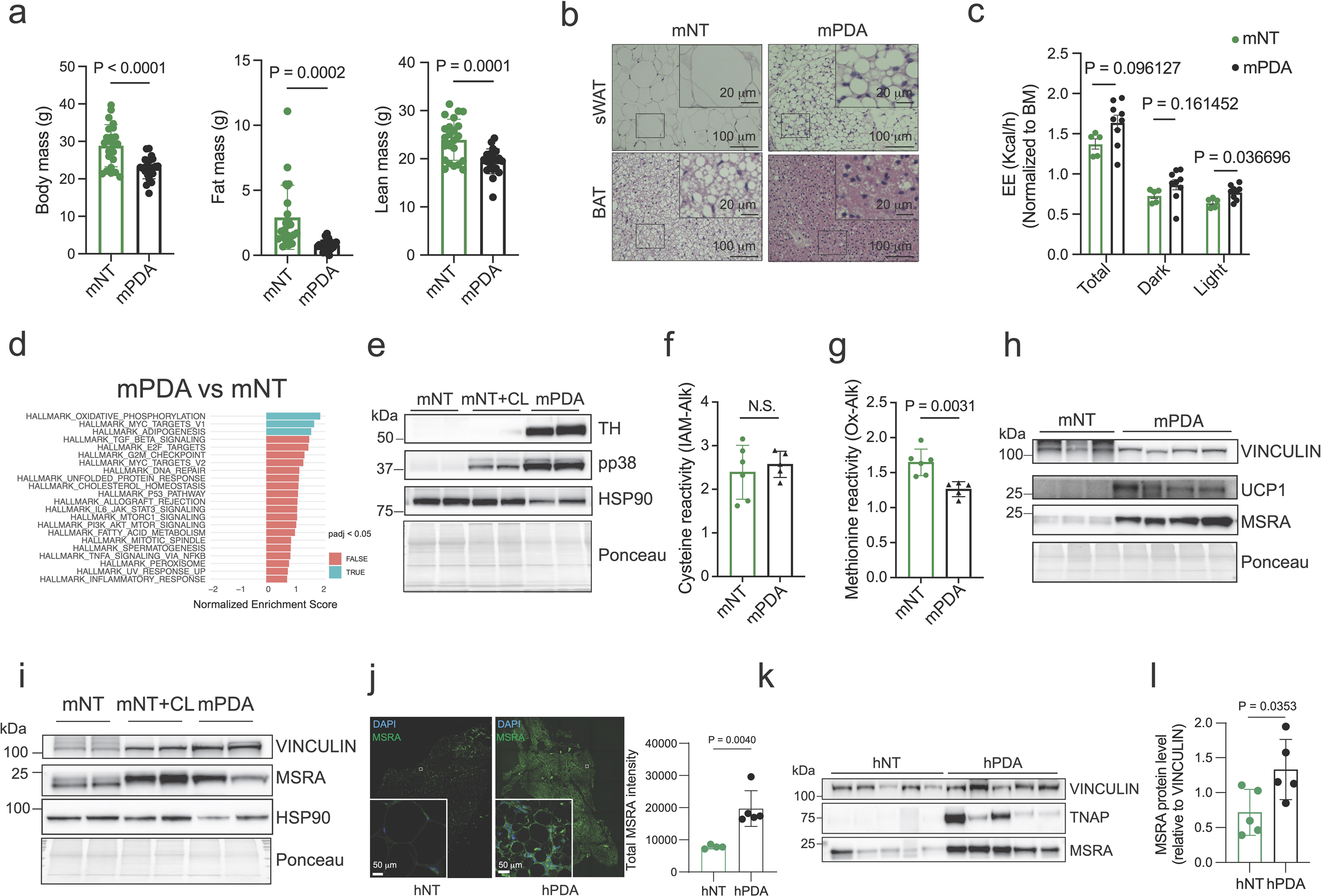
MSRA is markedly induced in white adipose tissues during cancer induced cachexia. **a**, Body, fat, and lean mass of KPC mice with tumors 7-9 mm in diameter (mPDA), compared to age-matched (6-8 months old) non-tumor (mNT) controls. **b**, Representative Hematoxylin and Eosin (H&E) staining of brown (BAT) and subcutaneous white (sWAT) adipose tissue of KPC mice with tumors 7-9 mm in diameter (mPDA), compared to age-matched non-tumor controls (mNT). **c**, Energy expenditure (EE) of KPC mice (mPDA) compared to age-matched-controls (mNT), normalized to body mass (BM). **d**, Gene Set Enrichment Analysis of the global proteome of sWAT from KPC mice (mPDA) compared to age-matched non-tumor controls (mNT). **e**, Immunoblot analysis of sWAT from KPC mice (mPDA) compared to age-matched non-tumor controls (mNT), and mice treated with 5 mg/kg β3-agonist CL-316243 (mNT+CL). **f**,**g**, Global cysteine oxidation analysis by iodoacetamide-alkyne (IAM-Alk) (**f**) and global methionine oxidation analysis by oxaziridine-alkyne (Ox-Alk) (**g**), comparing KPC mice (mPDA) and age-matched non-tumor controls (mNT). **h**, Immunoblot analysis of sWAT from KPC mice (mPDA) compared to age-matched non-tumor controls (mNT). **i**, Immunoblot analysis of sWAT from KPC mice (mPDA) compared to age-matched non-tumor controls (mNT), and mice treated with 5 mg/kg β3-agonist CL-316243 (mNT+CL). **j**, Immunohistochemical staining of MSRA in WAT from patients who underwent laparoscopic cholecystectomy (hNT) or were diagnosed with cachectic PDA (hPDA). **k**,**l**, Immunoblot analysis (**k**) and quantification (**l**) of WAT from patients who underwent laparoscopic cholecystectomy (hNT) or were diagnosed with cachectic PDA (hPDA). Error bars in this Fig. are means ± SDs. Student’s *t*-test was performed. N.S., not significant.

When non-shivering thermogenesis is induced by physiological stimuli, white adipocytes respond to the local release of norepinephrine by activating β3-adrenergic signaling, mitochondrial respiration, and UCP1 expression (Cannon and Nedergaard, 2004; Jimenez et al., 2003). In sWAT from cachectic KPC animals (Fig. 1e and Fig. S1g), we observed elevated levels of tyrosine hydroxylase (TH), the rate-limiting enzyme in the biosynthesis of catecholamine neurotransmitters and a surrogate for adipose sympathetic innervation (Chi et al., 2018). Moreover, global proteomic analysis of sWAT (Supplementary Table 1) revealed a marked upregulation of proteins involved in oxidative phosphorylation in both male (Fig. 1d) and female (Fig. S1h) KPC mice relative to controls. We also observed a similar induction of oxidative phosphorylation in non-tumor animals treated with the β3-AR agonist CL-316243 to mimic the physiological induction of WAT browning (Fig. S1i). Moreover, activated p38 kinase, a regulator of *Ucp1* transcription (Cao et al., 2004), is induced in sWAT of CL-316243-treated animals and tumor-bearing animals alike (Fig. 1e). Thus, the adipose atrophy that occurs in PDA-bearing mice is associated with a metabolic shift toward mitochondrial respiration consistent with a “browning” phenotype.

Since mitochondrial respiration generates reactive oxygen species (ROS) (Sena and Chandel, 2012), we expected that the intracellular oxidation state of WAT would be increased in PDA-bearing mice relative to controls. Thus, we surveyed the sWAT proteome for oxidative modifications such as lipid peroxide adduction (i.e, 4-HNE abundance) (Fig. S1j), cysteine oxidation (loss of IAM-Alk reactivity) (Fig. 1f), and methionine oxidation (loss of Ox-Alk reactivity) (Fig. 1g). Of these three parameters, only methionine reactivity was significantly perturbed in the tumor setting (Fig. 1g). These results suggested that oxidation of proteinaceous methionines may contribute to the pathobiology of adipose wasting during cancer cachexia.

### Methionine sulfoxide reductase A (MSRA) induction occurs upon both physiological and cachectic adipose browning

The sulfur atom of some methionine sidechains can act as a functionally relevant redox switch that toggles between its reduced and oxidized states (He et al., 2022; Kato et al., 2019). This toggling occurs because ROS can convert methionine residues to an oxidized form, methionine sulfoxide, which in turn can be enzymatically reduced back to methionine by methionine sulfoxide reductase A (MSRA) (Moskovitz et al., 1995). Interestingly, this oxidoreductase is strongly upregulated in both the sWAT (Fig. 1h) and eWAT (Fig. S1k) of tumor-bearing KPC mice. MSRA expression is also elevated, though more modestly, in the BAT compared to WAT depots of healthy mice (Fig. S1l). In addition, it can be robustly induced in the WAT of healthy mice by treatment with the β3-AR selective agonist CL-316243 (Fig. 1i and Fig. S1m), indicating that MSRA expression in adipocytes is responsive to sympathetic stimulation.

To ascertain whether similar metabolic changes occur during human cachexia, we examined a panel of visceral adipose tissues procured from i) non-tumor control patients who had undergone laparoscopic cholecystectomy (hNT) and (ii) PDA patients with cachexia (>5% weight loss, hPDA) (Supplementary Table 2). PDA patients displayed a striking decrease in adipocyte diameter (Fig. S1n) coupled with increased UCP1 and mitochondrial TNAP expression (Fig. S1o), consistent with activation of adipose browning. Increased MSRA expression (Figs. 1j-1l, Fig. S1p) and activating phosphorylation of the lipolytic enzyme HSL (Fig. S1p) were also observed in cachectic PDA patients. Thus, MSRA expression is upregulated in the white adipose tissue of both PDA patients and PDA-bearing mice in association with the metabolic shift toward mitochondrial respiration that occurs during WAT browning.

### Iron uptake stabilizes MSRA protein and induces adipose browning during cancer cachexia

The brown color of thermogenic adipocytes derives from the high-density iron-porphyrin complexes of their mitochondrial electron transport chains (Enerbäck, 2009). To increase oxidative metabolism in brown and beige adipocytes, β3-AR signaling promotes iron uptake to support the higher demand for these iron-bound mitochondrial enzymes (Yook et al., 2021). Thus, the transferrin receptor (TFR1) is upregulated during adipose browning, and disrupting TFR1 expression can impair both mitochondrial biogenesis and the thermogenic functions of brown/beige adipocytes(Yook et al., 2021). Interestingly, we observed markedly increased TFR1 levels in the WAT of both KPC mice (mPDA, Fig. S2a) and cachectic PDA patients (hPDA, Fig. S2b). Moreover, inductively coupled plasma mass spectrometry (ICP-MS) revealed that iron levels are significantly elevated in the adipose depots of tumor-bearing mice (Fig. S2c) and correspondingly reduced in their sera (Fig. S2d). These observations suggest that iron influx is not only essential for physiological adipocyte thermogenesis (Yook et al., 2021) but also facilitates adipose browning during cancer cachexia.

To determine whether iron is required for MSRA induction during physiological browning, we co-administered healthy mice with CL-316243 and the iron chelator deferiprone (DFP). As expected (Yook et al., 2021), iron chelation mitigates the browning effects of CL-316243 at the histological level (Fig. S2e). In addition, the robust induction of MSRA expression that occurs in these cells upon β3-AR activation – and coincides with both TFR1 and UCP1 upregulation – is reversed by DFP treatment (Fig. 2a). Conversely, expansion of the labile iron pool by CRISPR-mediated depletion of the iron storage protein ferritin heavy chain (FTH1) (Galy et al., 2023) allows post-transcriptional MSRA induction to occur in WAT adipocytes even in the absence of adrenergic stimulation (Figs. S2f and S2g). These results suggest that iron can directly elicit MSRA upregulation.

**Figure 2.**
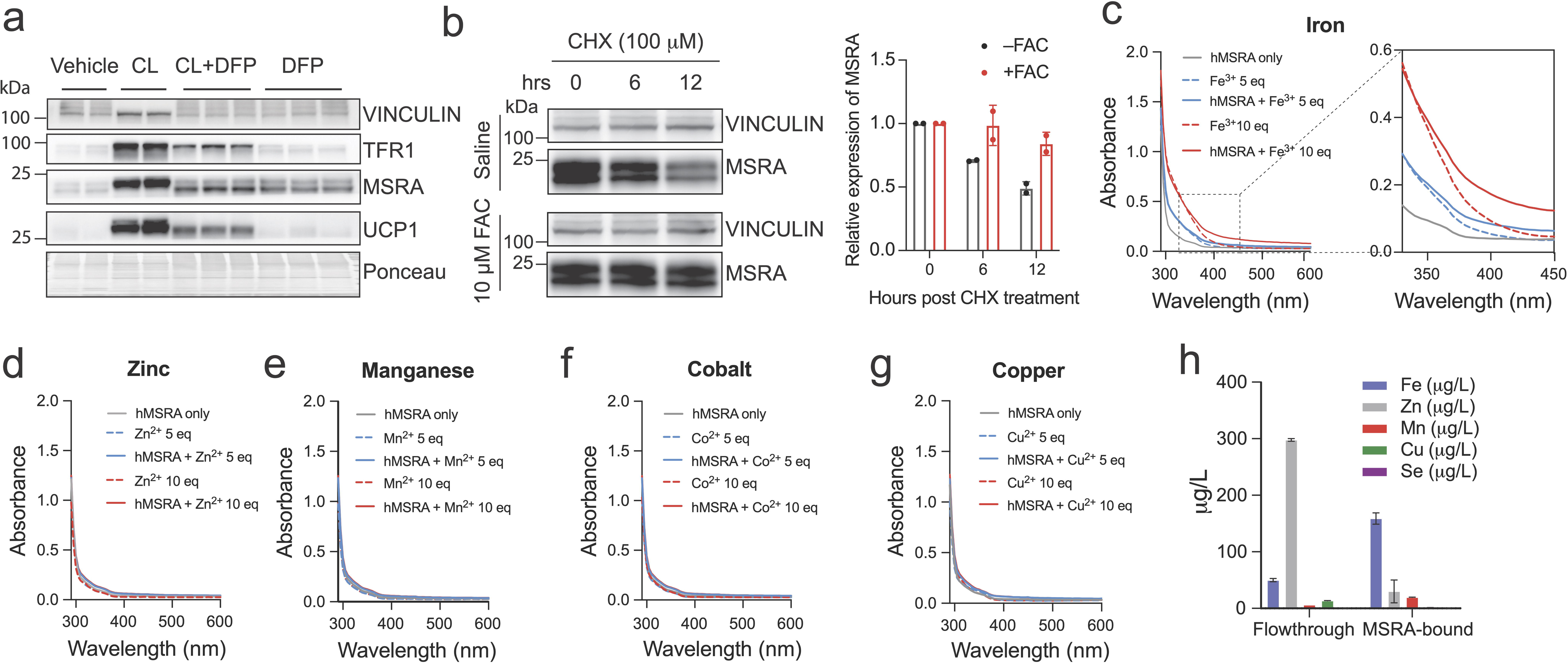
MSRA is stabilized upon iron binding. **a**, Immunoblot analysis of sWAT from mice treated with vehicle, 5 mg/kg β3-adrenergic agonist (CL-316243), 50 mg/kg deferiprone (DFP), or in combination (CL+DFP) for 6 days. **b**, Cycloheximide (CHX) pulse-chase to measure MSRA protein half-life in white adipocyte cultures treated with saline or ferric ammonium citrate (FAC). Right, quantification of MSRA relative to VINCULIN. **c**-**g**, Absorbance properties of recombinant human MSRA in the presence of different elements. Inset, zoomed-in spectra of MSRA in the presence of ferric ammonium citrate (FAC). 5eq, five molar equivalents in concentration. 10eq, ten molar equivalents in concentration. **h,** Inductively coupled plasma mass spectrometry (ICP-MS) is used to measure intracellular trace elements that are enriched in MSRA immunoprecipitants.

In mammalian cells, iron homeostasis is partly maintained by the iron regulatory proteins IRP1 and IRP2, which coordinate the expression of various iron metabolism genes at the post-transcriptional level by recognizing iron-responsive elements (IREs) within the 5’ and 3’-UTRs of their mRNA transcripts (Campillos et al., 2010; Galy et al., 2023). Although iron-dependent MSRA induction occurs post-transcriptionally (Figs. S2f and S2g), an *in silico* search (Campillos et al., 2010) failed to identify candidate IREs in the UTR sequences of mouse or human MSRA. We therefore asked whether iron can instead upregulate MSRA expression post-translationally by enhancing the stability of its protein product. Indeed, pulse-chase experiments revealed that the half-life of MSRA protein is markedly prolonged when adipocyte cultures are supplemented with iron in the form of ferric ammonium citrate (FAC) (Fig. 2b).

### Iron promotes the formation of an active MSRA oligomer

To determine whether this stabilization of MSRA involves a direct physical interaction with iron, we first analyzed its metal-binding properties *in vitro* and *in vivo*. Of note, the absorbance spectrum of purified recombinant MSRA was markedly shifted by exposure to ferric ions (Fe^3+^) but not to other metals, including Zn^2+^, Co^2+^, Mn^2+^, and Cu^2+^ (Figs. 2c-2g). Using ICP-MS to identify trace elements bound to endogenous MSRA protein, we further confirmed that iron content is substantially enriched in MSRA immunoprecipitants relative to other metals of equal or lower cellular abundance (Fig. 2h) and that this enrichment is further increased in cells supplemented with FAC (Fig. S2h). Together, these data suggested that MSRA is a metalloprotein with selective iron-binding capacity.

To ascertain how MSRA interacts with iron, we next examined its amino acid sequence using the metal ion-binding site prediction algorithm MIB2 (Lin et al., 2016). This analysis suggested that residues E203 and H206, which are highly conserved in MSRA orthologs across the evolutionary spectrum (Fig. 3a), have iron-binding potential (Fig. S3a). Moreover, site-directed mutagenesis of these two conserved residues (<u>E</u>DY<u>H</u> → <u>A</u>DY<u>A</u>), but not the MSRA myristoylation site (G22A) or active-site cysteine (C72S), significantly reduced MSRA protein expression levels (Fig. 3b). The EXXH motif of MSRA bears some resemblance to well-defined Fe-binding centers of several non-heme proteins that coordinate iron through the carboxylate and imidazole groups, respectively, of their conserved glutamic acid and histidine residues (Andrews, 2010). In particular, most members of the ferritin-like protein superfamily, including the f3 subunit of class I ribonucleotide reductases (RNRs), possess two conserved EXXH motifs that cooperate to form a diiron center comprised of two oxygen-bridged iron atoms (Banerjee et al., 2022; Ruskoski and Boal, 2021). For example, the two EXXH motifs of RNR, although widely separated within its primary amino acid sequence (Fig. S3a), converge in space to coordinate its diiron center (Figure 3c) (Banerjee et al., 2022; Ruskoski and Boal, 2021). Interestingly, the electrostatic environment surrounding the predicted iron-binding EXXH motif of MSRA bears a striking similarity to a half-site of the RNR diiron center (Figure 3c), we therefore, reasoned that MSRA oligomerization might allow for the assembly of a full diiron center, which in turn might enhance the stability, and even the enzymatic activity, of MSRA. Indeed, native PAGE analysis of adipocyte extracts revealed that MSRA exists in two conformations (Fig. 3d), of which the lower molecular weight species (presumed monomer) is stable and insensitive to exogenous iron. In contrast, the higher molecular weight form (presumed dimer) is labile but can be readily stabilized by iron supplementation (Fig. 3d). Moreover, iron chelation with DFP decreases expression of the monomeric and, to a greater degree, the dimeric form of MSRA (Fig. S3b). Interestingly, *in silico* analyses of mouse *apo* MSRA (PDB code: 6AGV), using either AlphaFold2-based multimer prediction (Liu et al., 2023) or COMPACK crystal packing similarity (Chisholm and Motherwell, 2005), predict that MSRA can assume a weak dimeric conformation mediated by hydrophobic interactions between residues E40, P43, T45, and D204 of each protomer (Fig. 3e). Moreover, in this conformation the EXXH motifs of both protomers are spatially juxtaposed, potentially allowing for coordination of an iron center which, in turn, could further stabilize the dimeric complex (Fig. 3e). In support of this notion, *in vivo* assembly of the dimeric MSRA species is largely abolished by either combined (AXXA) or individual (E203A and H206A) mutations of the EXXH motif (Fig. 3f). Moreover, size exclusion chromatography of recombinant MSRA shows that iron can enhance *in vitro* formation of a higher molecular weight species by wildtype MSRA but not by the MSRA-AXXA mutant (Fig. S3c). Importantly, the higher molecular weight species of MSRA corresponds to the enzymatically active reductase, as it can reduce methionine sulfoxide to methionine in a dose-dependent manner (Figs. S3d-S3f) and to the same degree as equimolar unfractionated MSRA-WT (Fig. S3g). In contrast, the MSRA-AXXA mutant exhibits no reductase activity (Figs. 3g and 3h). Thus, the iron influx induced by β3-AR stimulation of adipose tissues likely promotes dimerization, stabilization, and enzymatic activation of MSRA through the formation of an EXXH-coordinated diiron center.

**Figure 3.**
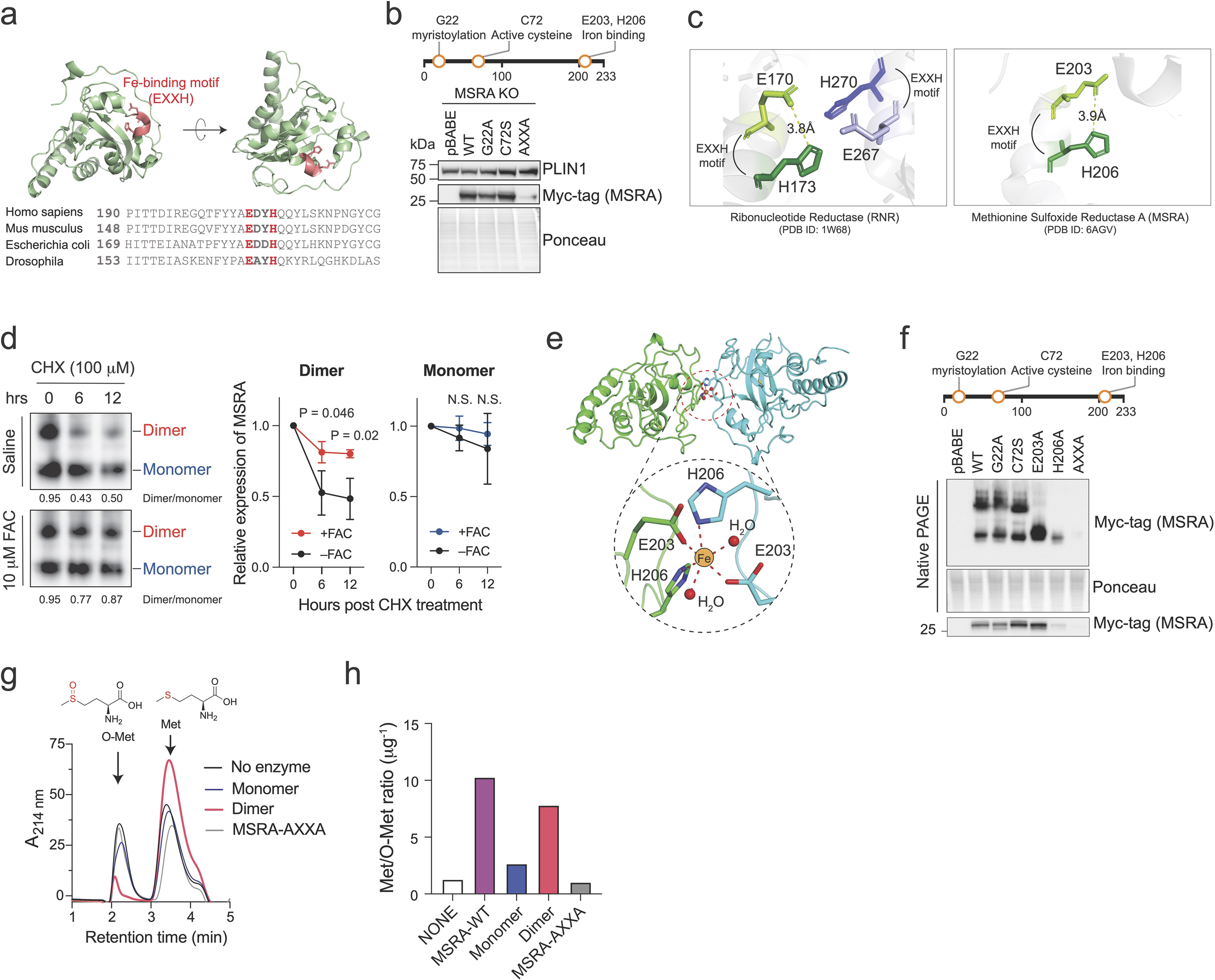
Iron binding promotes an active MSRA dimer. **a**, *Mus Musculus* MSRA crystal structure and sequence alignment. Red, putative iron (Fe) binding motif. **b**, Immunoblot analysis of wildtype (WT) and various point mutants of MSRA in white adipocyte cultures. G22A, myristoylation mutant; C72S, enzyme-dead mutant; AXXA, E203A-H206A double mutant; pBABE, empty vector control. **c**, Structural comparison of Ribonucleotide Reductase and MSRA iron-binding sites. **d**, Native PAGE analysis of MSRA protein half-life in white adipocytes upon cycloheximide (CHX) treatment, in the presence of saline or 10 μM FAC. Right, quantification. **e**, Predicted iron binding model of MSRA dimer. **f**, Native and SDS PAGE analysis of wildtype (WT) and point mutants of MSRA. G22A, myristoylation mutant; C72S, enzyme-dead mutant; E203A/H206A, single point mutants of iron-binding motifs; AXXA, E203A-H206A double mutant; pBABE, empty vector control. **g**,**h**, Enzymatic activity of gel filtration-purified MSRA monomer and dimer (**g**) are quantified (**h**) using the area ratio of reduced methionine (Met) and oxidized methionine (O-Met). Error bars in this Fig. are means ± SDs. Student’s *t*-test was performed. N.S., not significant.

### MSRA is required for **β**3-AR-induced PKA activation in adipocytes

To ascertain whether β3-AR signaling in adipose tissue is dependent on MSRA, we examined the phosphorylation of protein kinase A (PKA) substrates, a key mediator of the β3-AR-cyclic-AMP signaling axis (Granneman and Moore, 2008). As expected, global PKA substrate phosphorylation is upregulated in the WAT of non-tumor mice treated with CL-316243 relative to untreated controls (Fig. 4a; compare mNT+CL to mNT). Notably, a more pronounced induction is observed in the cachectic WAT of untreated PDA mice (Fig. 4a; compare mNT to mPDA), as well as PDA patients (Fig. 4b and Fig. S4a). Given its established role as a reductase of methionine sulfoxides(Moskovitz et al., 1995), we next asked if MSRA regulates the redox status of the PKA catalytic subunit. To this end, we used oxaziridine-alkyne (Ox-Alk) and iodoacetamide-alkyne (IAM-Alk) as chemical probes that selectively react with methionine and cysteine residues, respectively, but not with their oxidized derivatives (He et al., 2022) (Fig. S4b). Although IAM-Alk reactivity on the cysteine residues of PKA is comparable in WAT derived from PDA patients and their controls, the Ox-Alk reactivity of PKA methionines is significantly higher in the WAT of PDA patients (Fig. 4c) and PDA-bearing mice (Fig. 4d).

**Figure 4.**
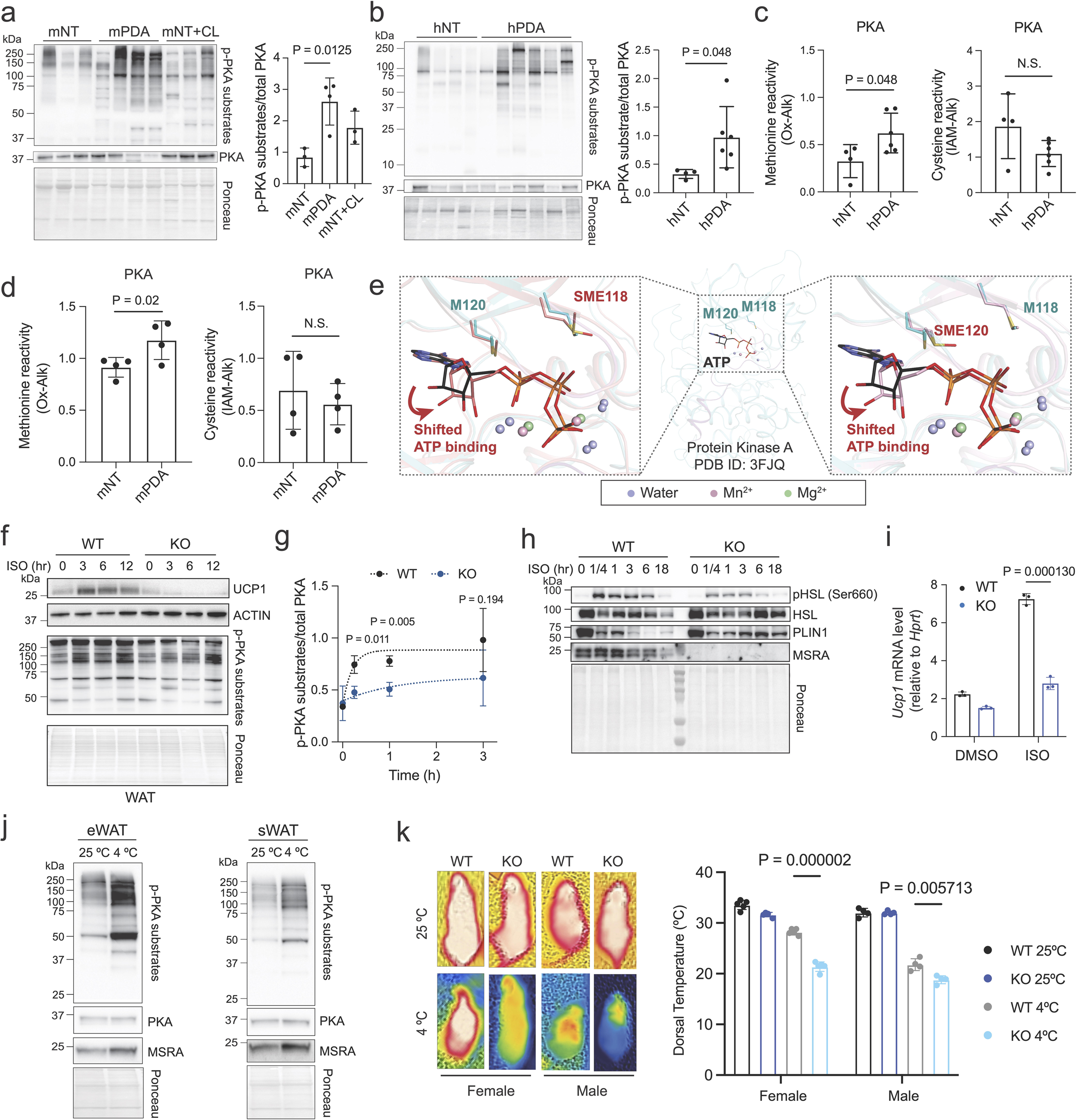
MSRA is required for PKA activation and adipose browning. **a**, Immunoblot analysis of sWAT from non-tumor controls (mNT), KPC mice (mPDA), and mice treated with 5 mg/kg β3-agonist CL-316243 (mNT+CL). Right, the level of PKA substrate phosphorylation is quantified relative to the total PKA expression in each setting. **b**, Immunoblot analysis of WAT from patients who underwent laparoscopic cholecystectomy (hNT) or were diagnosed with cachectic PDA (hPDA). Right, the level of PKA substrate phosphorylation is quantified relative to the total PKA expression in each setting. **c**,**d** PKA methionine oxidation status by oxaziridine-alkyne (Ox-Alk) and cysteine oxidation status by iodoacetamide-alkyne (IAM-Alk) analysis in human (**c**) and mouse (**d**) WAT. **e**, Structure overlay of *holo* WT PKA (cyan, PDB code 3FJQ) and molecular dynamic (MD) simulated models (Schrödinger Suite) of oxidized methionine (SME) forms of PKA at position 118 (SME118, red) and 120 (SME120, red). The cofactor ATP is shown as stick models for WT PKA (in black) and MD simulated models when M118 is oxidized (SME118, in salmon), and when M120 is oxidized (SME120, in pink). **f**,**g**, Immunoblot analyses of white (WAT) adipocyte cultures upon stimulation with 100 μM pan-β-adrenergic receptor agonist, isoproterenol (ISO) over the indicated number of hours (**f**). The level of PKA substrate phosphorylation is quantified relative to the total PKA expression in at each time point (**g**). **h**, Immunoblot analyses of white adipocyte cultures upon stimulation with 100 μM pan-β-adrenergic receptor agonist, isoproterenol (ISO) over 18 hours. **i**, qRT-PCR analysis of white adipocyte cultures upon treatment with DMSO vehicle or 100 μM pan-β-adrenergic receptor agonist, isoproterenol (ISO) for 24 hours. **j**, Immunoblot analysis of WAT tissues of male mice housed in room temperature or in 4°C for 6 hours. **k**, Surface body temperature, calculated as the average of the highest 50% of the region of interest, of adult mice before and after 6 hours of acute cold exposure. WT, *MsrA^+/+^*; KO, *MsrA^−/−^*. Error bars in this Fig. are means ± SDs. Student’s *t*-test was performed. N.S., not significant.

Of the six methionines in the catalytic subunit of PKA, two residues (M118 and M120) are situated near its ATP-binding site and both are highly solvent-exposed (Fig. 4e). A structural overlay of *holo* ATP-Mn^2+^-bound PKA (PDB code: 3FJQ) with that of ADP-bound PKA (PDB code: 1JBP) reveals negligible positional variation in the AMP moieties of ADP and ATP between the two structures (Fig. S4c). Moreover, ATP and the metal ions of *holo* PKA are tightly confined in space by the active site residues (Thompson et al., 2009), presumably to optimize catalysis (Fig. S4c). However, molecular dynamics simulations using Schrödinger software revealed striking conformational changes of *holo* PKA when residues M118 and M120 are oxidized either individually (Fig. 3e) or in combination (Fig. S4d). In either case, methionine oxidation displaces the adenine and ribose groups by 0.7 Å and 1.4 Å, respectively (Fig. 4e and Fig. S4d). In addition, oxidation of either M118 or M120 causes local steric clashes with neighboring residues as determined by Phenix refinement (Afonine et al., 2012) of the *holo* enzyme x-ray diffraction data (PDB code: 3FJQ); moreover, this effect is even more pronounced when both methionine residues are oxidized (Fig. S4e). These *in silico* findings suggest that oxidation of M118 and/or M120 can impair PKA enzymatic function by disrupting ATP-binding.

To assess the functional relationship between MSRA and PKA experimentally, we first examined adipose tissues of *MsrA*^−/−^ mice, which mature to adulthood with no apparent phenotype in the absence of oxidative stress (Moskovitz et al., 2001). The BAT and WAT of *MsrA*-null animals are histologically indistinguishable from those of wildtype mice (Fig. S4f), and the BAT-specific expression of UCP1 is unaffected by *MsrA* nullizygosity (Fig. S4g). These results indicate that *MsrA* is not required for adipocyte development *in vivo*. Likewise, *ex vivo* differentiation of cultured pre-adipocytes from neonatal adipose tissues (Galmozzi et al., 2021) does not require *MsrA* (Fig. S4h). However, while the pan-βAR agonist isoproterenol induces global PKA substrate phosphorylation and UCP1 expression in white adipocyte cultures (Figs. 4f and 4g), this effect is impaired in *MsrA*-null cells, despite comparable PKA levels (Fig. S4i). *MsrA* loss also dampens other downstream consequences of β3-AR/PKA signaling, such as activating phosphorylation of hormone-sensitive lipase (HSL) (Fig. 4h and Fig. S4j) and transcriptional induction of the thermogenic gene *Ucp1* (Granneman and Moore, 2008) (Fig. 4i). Consistent with a role for MSRA in non-shivering thermogenesis, its expression is induced in WAT upon acute cold exposure (Fig. 4j), and the ability of *MsrA*^−/−^ mice to maintain body temperature in response to cold is significantly impaired (Fig. 4k). Thus, MSRA is required for β3-AR-dependent induction of PKA enzymatic activity and the thermogenic program of white adipocytes.

### MSRA is essential for PDA-associated adipose atrophy and cachexia

To determine if the thermogenic functions of MSRA are also required for adipose remodeling in response to tumor development, we interbred KPC and *MsrA*^−/−^ mice to generate KPCM compound-mutant (*Kras^G12D^; p53^R172H^; PdxCre; MsrA^−/−^*) animals. Comparing KPC and KPCM mice with similar tumor burdens revealed that body mass attrition is significantly mitigated by the absence of *MsrA*, both during early (tumor 5-8 mm diameter) (Fig. S5a) and late (tumor 9-10 mm diameter) (Fig. 5a) stages of disease. The rate of body mass depletion was also reduced in KPCM mice when measured over a 4-week period after initial detection of a palpable tumor (Fig. S5b). *MsrA* deletion also preserved adipose tissue volume (Fig. 5b) and the unilocular features of sWAT in mice bearing early-stage tumors (Fig. 5c). Interestingly, although MSRA expression is not induced in the skeletal muscles of KPC mice (Fig. S5c), the absence of *MsrA* did preserve lean mass (Fig. S5d) and skeletal myocyte diameter (Fig. S5e) in KPCM mice, consistent with the notion that adipose atrophy precedes muscle wasting during cancer cachexia (Kir et al., 2014; Neves et al., 2016; Petruzzelli et al., 2014; Zimmers et al., 2016). No apparent effects of *MsrA* loss were observed on primary tumor burden (Extended Data Fig. 5f), the macroscopic or histological appearance of other organs (Fig. S5g), food intake (Fig. S5h), sWAT sympathetic tone (Fig. S5i), or sWAT β3-AR expression (Fig. S5j).

**Figure 5.**
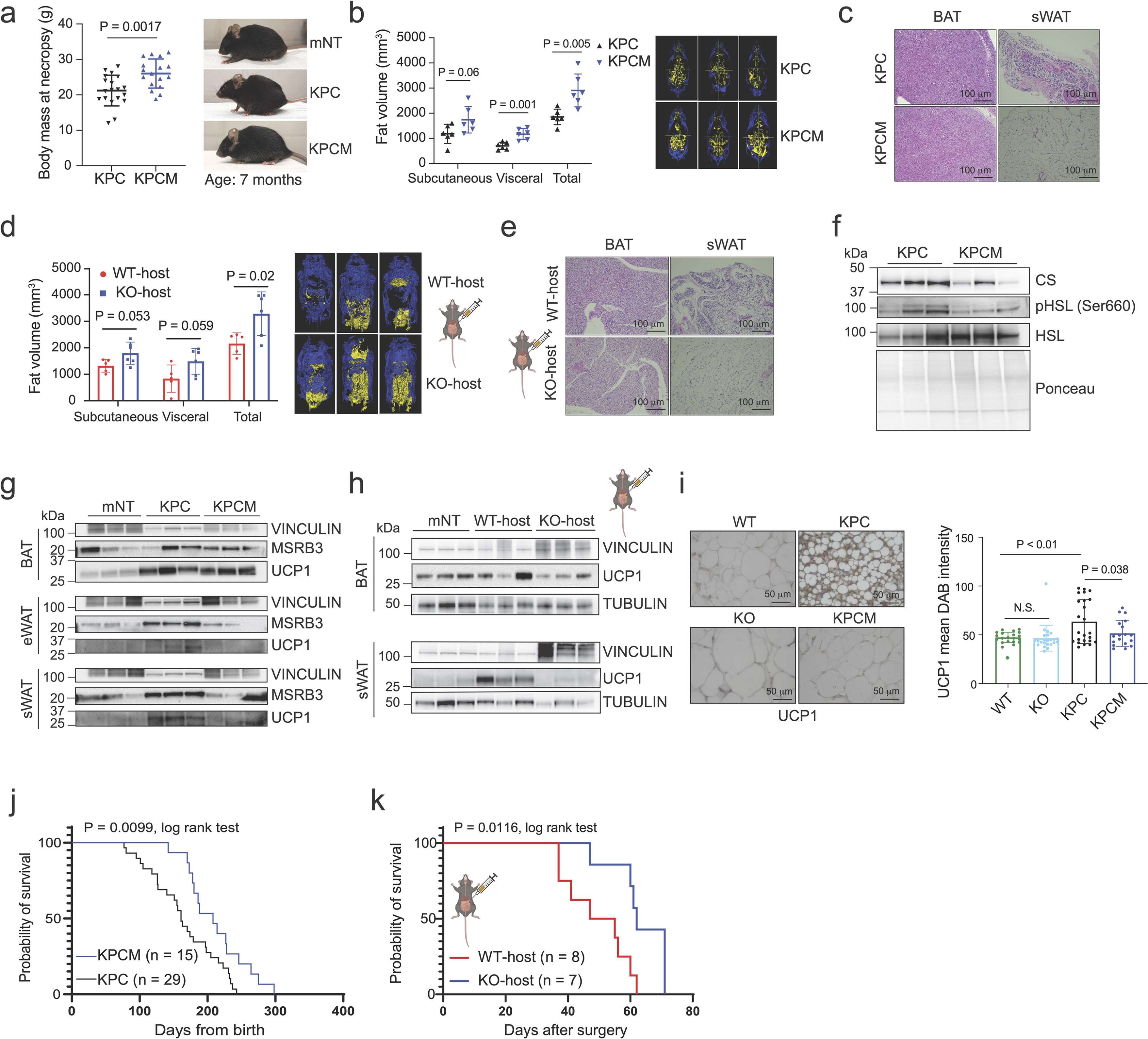
MSRA is required for PDA-associated adipose atrophy and cachexia. **a**, Body mass of pancreatic tumor-bearing mice. KPC, *Kras^G12D^;p53^R172H^;PdxCre;MsrA^+/+^*(n = 18; 5 males and 13 females); KPCM, *Kras^G12D^;p53^R172H^;PdxCre;MsrA^−/−^*(n = 17; 7 males and 10 females). **b**, Fat volume analysis by magnetic resonance imaging (MRI). KPC, n = 6 (4 males and 2 females); KPCM, n = 6 (3 males and 3 females). **c**, Representative Hematoxylin and Eosin (H&E) staining of brown (BAT) and subcutaneous white (sWAT) adipose tissue of mice with pancreatic tumors 7-9 mm in diameter. **d**, Fat volume analysis by MRI of male mice 5 weeks after orthotopic pancreatic cancer cell injection. WT-Host, *MsrA^+/+^*; KO-host, *MsrA^−/−^*. **e**, Representative Hematoxylin and Eosin (H&E) staining of BAT and sWAT of mice 5 weeks after orthotopic pancreatic cancer cell injection. **f**-**h**, Immunoblot analysis of BAT, eWAT, and sWAT from autochthonous PDA mice (**f and g**) and mice 5 weeks after orthotopic pancreatic cancer cell injection (**h**). mNT, Non-tumor control. **i**, Immunohistochemical analysis of UCP1 expression in sWAT of tumor-bearing mice compared to age-matched non-tumor controls. Right, quantification of UCP1 intensity. Values plotted are average from 5 fields of view from each animal. WT, non-tumor *MsrA^+/+^*; KO, non-tumor *MsrA^−/−^.* **j**,**k**, Overall survival of autochthonous (**j**) and orthotopic transplant (**k**) mouse models of PDA. KPC (n = 29; 13 males and 16 females); KPCM (n = 15; 4 males and 11 females); WT-host, *MsrA^+/+^* (n = 8 males); KO-host, *MsrA^−/−^*(n = 7 males). Log rank test was performed. Error bars in this Fig. are means ± SDs. Student’s *t*-test was performed unless stated otherwise. N.S., not significant.

To assess the possibility of cancer-cell intrinsic MSRA functions in PDA cachexia (He et al., 2022), we further asked whether PDA cells induce adipose and muscle atrophy to a different degree when orthotopically transplanted into hosts that are either wildtype (WT-host) or nullizygous (KO-host) for *MsrA*. Consistent with the KPC model, male mice transplanted orthotopically with PDA cells lose body mass during tumor progression (Extended Data Fig. 6a) at a rate that correlates inversely with overall survival (Extended Data Fig. 6b). Consistent with adipose atrophy as an early event in cancer cachexia (Abdullahi and Jeschke, 2016; Arner and Langin, 2014; Seelaender and Batista, 2014), loss of adipose tissue mass coincides with gain of tumor mass, and both occur before changes in lean mass are detectable (Extended Data Fig. 6c). At 4-5 weeks post-surgery, a time when adipose depletion is robust (Extended Data Fig. 6c), the transcriptome of sWAT (Supplementary Table 3) from tumor-bearing mice is strongly enriched for oxidative phosphorylation (Fig. S6d), as well as markers of adipose browning such as *Elovl3*, *Ucp1*, and *Dio2* (Fig. S6e). Having established this timeline, we then performed orthotopic injections of PDA cells into cohorts of age-matched *MsrA^+/+^* (WT-host) and *MsrA^−/−^*(KO-host) male hosts. As seen in the autochthonous model (Figs. 5a-4c), *MsrA* deletion in the host mitigates adipose (Fig. 5d) and muscle (Extended Data Fig. 6f) loss in tumor-bearing mice five weeks after tumor inoculation, while having no effect on primary tumor burden (Extended Data Fig. 6g) or the incidence of metastasis (Fig. S6h). Histologically, *MsrA* loss in the host mitigates development of multilocular morphology by WAT cells of tumor-bearing mice (Fig. 5e). At the molecular level, mitochondrial content as measured by citrate synthase levels, phosphorylating activation of HSL (Fig. 5f), and the induction of UCP1 are all impaired by MSRA deletion (Figs. 5g-i). These effects cannot be attributed to changes in sympathetic innervation (Fig. S6i) or total iron content (Fig. S6j), as these parameters are comparable in wildtype and *MsrA*-deficient adipose tissues. Also, *MsrA* loss does not lead to compensatory changes in the expression of the methionine sulfoxide reductase B proteins (MSRB2 and MSRB3) (Figs. S6k and S6l).

Since loss of adipose tissue is linked to reduced survival in PDA patients (Fouladiun et al., 2005; Murphy et al., 2010), we asked whether MSRA-dependent adipose depletion affects survival outcomes in PDA-bearing mice. Of note, *MsrA* deletion significantly prolonged overall survival in both the autochthonous (Fig. 5j) and orthotopic transplant (Fig. 5k) models of PDA. These improvements in survival upon *MsrA* deletion are not associated with either a decrease in primary tumor burden (Fig. S7a) or the incidence of metastasis (Fig. S7b). Taken together, our findings identify MSRA as a key regulator of adipose remodeling that is essential for thermogenic browning and the initiation of cachectic phenotypes during pancreatic tumorigenesis.

## DISCUSSION

Cachexia is a debilitating disorder that plagues nearly half of cancer patients. Its biological origins remain poorly understood and there are still no effective therapies to alleviate its lethal impact (Baracos et al., 2018). In the context of PDA, the browning and depletion of white adipose tissues (WAT) begins early in the course of disease, even before clinical diagnosis of PDA(Sah et al., 2019). Therefore, to elucidate mechanisms that activate WAT browning and atrophy, we examined specimens from cachectic PDA patients and employed both autochthonous and orthotopic transplant models of PDA. These studies revealed that iron metabolism, and consequently the redox state of proteinaceous methionine residues, is specifically modified during WAT browning. Moreover, we show that MSRA, an enzyme that can reduce oxidized methionine residues, is essential for WAT browning in both physiological and cachectic settings.

A recent study revealed that physiological thermogenesis in browning adipocytes is dependent on an expansion of the labile iron pool (Yook et al., 2021). Here, we show that iron influx also occurs in browning adipocytes during cancer cachexia in both PDA patients and mouse PDA models. While increased iron content would support the synthesis of the iron-sulfur clusters needed to power mitochondrial respiration (Yook et al., 2021), in principle, it could also promote ROS production through the Fenton reaction (Lloyd et al., 1997), rendering proteins more susceptible to oxidative modifications during browning. Indeed, we observed that the methionine proteome of PDA WAT is more oxidized globally. Since methionine oxidation converts a hydrophobic residue to a polar one, this modification is thought to destabilize proteins (Brot and Weissbach, 1983) or exacerbate protein misfolding and aggregation (Younan et al., 2012). In addition, however, we and others have shown that certain methionine residues of select proteins can act as functional redox switches that govern particular cellular processes (He et al., 2022; Kato et al., 2019). Thus, the sulfur atom of these residues can toggle between its reduced (methionine) and oxidized (methionine sulfoxide) states, the balance of which is controlled by changes in MSRA activity. Here we demonstrate that iron influx and MSRA induction are both essential for WAT browning, and that the enzymatic activity of MSRA requires direct interaction with iron. More specifically, residues E203 and H206 of human MSRA comprise a phylogenetically conserved EXXH iron-binding motif, similar to those common among proteins of the ferritin superfamily (Andrews, 2010). Ferritin-like proteins typically harbor two EXXH motifs which cooperate to form a single iron-binding center. In contrast, MSRA, with its solitary motif, will only coordinate iron upon multimerization, when a complete iron-binding center can be formed by spatially juxtaposed EXXH motifs from two MSRA polypeptides. Since multimerization promotes both its stability and enzymatic activity, MSRA is ideally suited to function in the iron-rich milieu of browning WAT cells.

In general, adrenergic signaling, including the induction of adipose browning by β3-AR stimulation, is dependent on PKA phosphorylation of its appropriate substrates. Here we show that MSRA maintains PKA in an active state during WAT browning likely by erasing oxidation of two solvent-exposed methionines near its ATP-binding site (residues M118 and M120). Consistent with our findings, the M120A point mutation has been shown to suppress PKA enzymatic activity by 60% (Morgan et al., 2008). MSRA is often viewed as a housekeeping enzyme that acts constitutively to maintain redox homeostasis across the proteome in multiple cell types (Moskovitz et al., 2001). However, our work reveals that the iron-dependent multimerization and enzymatic activation of MSRA allows it to function more robustly in adipose cells that experience iron influxes during either physiological or cachectic adipose browning. Moreover, its ability to reverse oxidation of critical methionines, such as PKA residues M118 and M120, may render MSRA essential in these particular settings – and also explain why MSRA is required for adipose browning but not more broadly in normal animal development (Moskovitz et al., 2001).

Understanding the differences between physiological browning, a reversible process associated with metabolic health (Abdullahi and Jeschke, 2016; Nedergaard and Cannon, 2014), and cachectic browning, which is irreversible even in the face of increasing caloric intake (Baracos et al., 2018), should facilitate development of clinical therapies to alleviate cancer cachexia. While PKA oxidation was examined in depth in the current study, global chemoproteomic profiling may also uncover additional MSRA effector proteins, including some that are differentially regulated in browning during thermogenesis versus cancer cachexia. In any case, here we have observed that genetic inactivation of MSRA is sufficient to prevent cachexia and thereby prolong the survival of PDA-bearing animals. Thus, the unique iron metabolism of browning adipocytes should be an effective and relatively nontoxic target for pharmacological suppression of cachexia in cancer patients.

## Acknowledgments

We are grateful to Huijin Feng for technical assistance, Minhee Lee for HPLC instrument training, and Nicolae Ciobu Zubenco for performing tail vein injections. We are grateful to Dr. Bruce Spiegelman for his guidance on adipose tissue analyses. We also thank other members of the Chio lab, as well as Drs. Richard Baer, Neel Shah, and Viraj Sangvhi for discussion. We thank J.R. Prigge and Ed Schmidt, Montana State University, for developing the anti-Nrf2 antibody. This work was performed with the support of the Herbert Irving Comprehensive Cancer Center (Columbia University Irving Medical Center) Proteomics, Flow Cytometry, Genomics and High Throughput Screening, Oncology Precision Therapeutics and Imaging Core, and Molecular Pathology (MPSR) Shared Resources, as well as the Mass Spectrometry Core Facility (Chemistry Department at Columbia University). AAV transfer plasmid cloning and packaging were performed by the Zuckerman Institute’s Genetic Access Tools and Virology Platforms (Columbia University). We thank Dr. Hanina Hibshoosh (MPSR) for coordinating the procurement of surgically resected specimens of adipose tissues, as well as Drs. Neel Shah for sharing resources for the synthesis of the oxaziridine probe. **Funding:** This work was supported by National Institute of Health (NIH) grants (R01-CA240654, 1R01CA267870, 1R01CA273023 to I.I.C.C., P30ES009089 to K.S, and R50CA210240 to P.G.), Pershing Square Sohn Research Alliance (to I.I.C.C.), Irma Hirchl Trust (to I.I.C.C). The University of Nebraska Medical Center’s Rapid Autopsy Program for Pancreas is supported by SPORE in Pancreatic Cancer (P50CA127297), Pancreatic Cancer Detection Consortium, (U01CA210240), NCI Cancer Center Support Grant (P30CA36727). The Herbert Irving Comprehensive Cancer Center at Columbia University is supported by P30-CA13696 and the Columbia University flow cytometry core is supported by P30CA36727.

## STAR METHODS

### RESOURCE AVAILABILITY

#### Lead contact

Further information and requests for resources and reagents should be directed to and will be fulfilled by the lead contact, Iok In Christine Chio.

#### Materials availability

All cell lines and plasmids generated can be obtained via a CUIMC materials transfer agreement (free of charge for noncommercial purposes).

#### Data and code availability

The mass spec datasets generated for these studies are available as supplemental data provided with this manuscript.

### EXPERIMENTAL MODEL AND SUBJECT DETAILS

#### Human Patient Samples

##### Rapid Autopsy Samples

Visceral adipose tissue samples were obtained from individuals previously diagnosed with pancreatic ductal adenocarcinoma through the University of Nebraska Medical Centre’s Rapid Autopsy Program (UNMC RAP) for Pancreas under IRB 091-01. Non-cancerous tissues were collected in a similar manner through the UNMC Normal Organ Recovery Program in collaboration with LiveOn Nebraska. To ensure specimen quality, organs were harvested within three hours post-mortem, and the samples were either flash-frozen in liquid nitrogen or immediately placed in formalin for fixation.

##### Columbia University samples

Visceral adipose tissue was freshly collected from human patients undergoing laparoscopic cholecystectomy or resection of pancreatic ductal adenocarcinoma at Columbia University Irving Medical Center in compliance with IRB-AAAU7458.

#### Animal Models

##### Genetically engineered mouse model of pancreatic cancer

*Trp53^+/LSL-R172H^*, *Kras^+/LSL-G12D^*, *Pdx1-Cre,* strains in C57Bl/6NJ background were interbred to (KPC)(Hingorani et al., 2005) mice. *MsrA* knockout mice in C57BL/6J were obtained from Jackson Laboratory (RRID:MMRRC_060602-JAX) and interbred with KPC to generate compound mutants obtain *Kras^+/LSL-G12D^; Trp53^+/LSL-R172H^ ; Pdx1-Cre*; *MsrA^−/−^* (KPCM).

##### Orthotopic allograft model of pancreatic cancer

Orthotopic engraftment of mouse pancreatic cancer cells was conducted as described (He et al., 2022). In brief, mice were anesthetized using isoflurane and subcutaneously administered 5 mg/kg Carprofen. 10^5^ cells were transplanted to the parenchyma of the pancreas. The abdominal wall was sutured with absorbable vicryl sutures (Ethicon Cat# J392H), and the skin was closed with wound clips (CellPoint Scientific Inc. Cat# 203-1000). C57Bl/6J mice were purchased from the Jackson Laboratory (RRID:IMSR_JAX:000664) for syngeneic orthotopic transplant experiments.

Tumor-bearing mice were subjected to high-contrast ultrasound imaging using the Vevo 2100 System with a MS250, 13–24 MHz scanhead (Visual Sonics, Inc, Amsterdam, NL) to monitor tumor volume intravitally. All animal experiments were conducted in accordance with procedures approved by the IACUC at Columbia University (AC-AABK554).

### METHODS DETAILS

#### Indirect calorimetry system and body composition

Upon detection of tumor mass or after surgical implantation of cancer cells, male and female mice were acclimated in metabolic chambers (Promethion Core, Sable Systems International) for 4 days before the start of the recordings. Mice were continuously recorded for 3 days, with measurements taken every 30 minutes, including food intake, water intake, and ambulatory activity (in X and Z axes) using rodent metabolic cages. Values were adjusted by body mass where mentioned. Energy expenditure was analyzed after normalization based on body mass. In addition, food intake was continuously determined by integrating weighing sensors fixed at the top of the cage, from which the food containers had been suspended, into the sealed cage environment. Body composition was measured intravitally by MRI (EchoMRI; Echo Medical Systems, Houston, TX).

#### Magnetic Resonance Imaging (MRI) to measure body fat

##### Imaging

The MRI experiments were carried out using a Bruker BioSpec 9.4 Tesla preclinical scanner equipped with the Paravision (PV.7.0.0) software platform (Bruker Corp., Billerica, MA). Mice were anesthetized with 1-2% isoflurane mixed with medical air and delivered via a nose cone. During the procedure, the isoflurane concentration was adjusted to maintain a respiratory rate of 40-80 breaths per minute, monitored with a respiration pillow connected to a monitoring system (SA Instruments, Stony Brook, NY). The mice were placed in a prone position on the animal bed inside a circular polarized birdcage coil with a 72 mm inner diameter. Images were acquired using the Rapid Acquisition with Relaxation Enhancement (RARE) sequence with a fat-water separation technique. The imaging parameters were set as follows: TR = 1448 ms, TE = 26 ms, and a resolution of 265 x 265 μm². Coronal images were obtained with a slice thickness of 1mm, consisting of 24 slices to cover the entire body of the mouse. A 140° angle between fat and water magnetization was used to produce out-of-phase images. Separate images of fat and water were generated, and a recombined fat-water image was reconstructed to provide a clear depiction of the mouse’s anatomy.

##### Data Analysis

The MRI data were imported into Analyze Direct 14.0 software. Initial preprocessing steps included contrast enhancement to improve the clarity of the fat tissue boundaries. Segmentation was carried out to differentiate and quantify subcutaneous and visceral fat. Using the thresholding tool, intensity thresholds were set to identify fat tissues based on their specific signal characteristics. The segmentation process was further refined using region-growing algorithms to ensure accurate delineation of fat boundaries. Manual adjustments were made where necessary to correct any misclassifications. Following segmentation, the volume of the identified fat compartments was calculated. The software’s volume rendering module was used to create 3D reconstructions of the fat distribution. Volume measurements were obtained by summing the voxel volumes of the segmented regions. The total fat volume, as well as the volumes of subcutaneous and visceral fat, were recorded for each subject.

#### Acute cold challenge

Adult male and female mice were divided into 25 °C and 4 °C groups. The temperature-challenging duration was 6 hours. The compact thermal camera (FLIR C5) was used to measure the dorsal temperatures of individual mice, and the collected images were processed using the FLIR thermal studio. The temperatures were measured before and after the cold exposure.

#### *In vivo* **β**3-AR agonist and iron chelator treatment

To study the effects of β3-adrenergic receptor activation and iron chelation, the β3-adrenergic receptor agonist CL-316243 disodium salt (Cayman Chemical, Cat#17499) was systemically administered by daily intraperitoneal injection (5 mg/kg in saline), and deferiprone (EDQM, Cat#Y0001976) was administered by oral gavage (50 mg/kg in water). All mice were dosed for 6 days daily.

#### Immunohistochemistry

Tissues were fixed in 10% neutral buffered formalin and embedded in paraffin. Sections were subjected to H&E staining as well as immunohistochemical staining. Antigen retrieval was done in 10 mM citrate buffer (pH 6). The following primary antibodies were used for immunohistochemical staining at the indicated dilution: mUCP1 (RRID: AB_2687530, 1:500), mβ3AR (RRID: AB_2636362, 1:100), mTH (RRID: AB_297840, 1:250), hUCP1 (RRID: AB_2241462, 1:100), hMSRA (RRID: AB_2682032, 1:50).

#### Adipocyte and myocyte diameter analysis

H&E-stained slides of adipose and skeletal muscle tissues were scanned using the Leica SCN400 at 40x, and the diameter of 100 cells was measured per slide using the QuPath software (v0.5.1). Data was analyzed using Prism 8 (GraphPad software).

#### Cell culture

HEK293T cell line was from ATCC (Cat# CRL-3216). Primary white adipocytes were isolated from newborn mice based on a published protocol (Galmozzi et al., 2021). In brief, white adipose depots were harvested from pups on postnatal day 1. Tissues were dissociated with 15 mg/mL collagenase I for 30 minutes at 37 °C at 1400 revolutions per minute in a Thermomixer. Digest is put through a 100 μm cell strainer and plated. When preadipocytes become confluent, differentiation was performed with 10% DMEM, 170 nM insulin, 1 μM Dexamethasone, 0.5 mM 3-isobuthyl-1-methylxanine, and 2 μM Rosiglitazone. Unless stated otherwise, all cells were cultured in DMEM (Gibco Cat#11995073), supplemented with 10% fetal bovine serum (FBS) (Corning Cat# 35-010-cv) and 1% PS. All cells were cultured at 37 °C with 5% CO_2_.

#### CRISPR/Cas9-mediated gene deletion

White preadipocytes were transduced with lentivirus expressing Cas9 and sgRNAs (lentiCRISPRV2, Addgene # 52961). Single sgRNAs were cloned by annealing two DNA oligos and T4 DNA ligation into a BsmB1-digested LRC2.1T as described (Shi et al., 2015). The design principle of sgRNA was based on previous reports: (1) sgRNAs with the predicted low off-target effect (Hsu et al., 2013) and (2) sgRNAs targeting the functional protein domain region (Shi et al., 2015), for which sequence information was retrieved from the NCBI Conserved Domains Database. gRNA sequences used to target mouse *Fth1* are: GGAGAGCGGGCTGAATGCAA (sense), and TTGCATTCAGCCCGCTCTCC (antisense). gRNA sequence against the *Rosa26* locus was used as a negative control: GAAGATGGGCGGGAGTCTTC.

#### Lentiviral and retroviral production and infection

LentiCRISPRV2 lentiviruses were produced in HEK293T cells co-expressing the packaging vectors (pPAX2, Addgene # 12260 and VSVG, Addgene # 12259), concentrated with LentiX concentrator (Clontech Cat# 631232), and resuspended with DMEM supplemented with 10% FBS and 1% PS at 10X concentration. Mouse pBabe (Addgene # 1764 and # 1767) retroviruses were produced in Phoenix-ECO cells (ATCC Cat# CRL-3214), concentrated with RetroX Concentrator (Clontech Cat# 631456), and resuspended in DMEM supplemented with 10% FBS and 1% PS at 10X concentration. One hundred thousand cells were plated and infected with 5X concentrated viruses and spinoculated at 600 g for 45 min at room temperature. One day after infection, cells were treated with 3 μg/mL puromycin (Sigma-Aldrich Cat # P9620) for selection.

#### Global proteomics LCMS

##### Sample Preparation

iST Sample Preparation kit (PreOmics, Germany) was used according to the manufacturer’s protocols. Briefly, 50 μL of Lysis buffer was added and heated at 95 °C for 10 min at 1000 rpm with agitation. After cooling the sample to room temperature, trypsin digestion buffer was added, and the sample was incubated at 37 °C for 2 h at 500 rpm with shaking. Sample clean-up and desalting were carried out in the iST cartridge using the recommended wash buffers. Peptides were eluted with elution buffer (2 × 100 μL) and then lyophilized by SpeedVac.

##### Spectrogram Library Establishment 2.2.1 High pH Reverse Phase Separation

The peptide mixture was re-dissolved in buffer A (buffer A: H_2_O, pH 10.0, adjusted with ammonium hydroxide), and then fractionated by high pH separation using nanoACQUITY UPLC system (Waters Corporation, MA, USA) connected to a reverse phase column (C18 column, 2.5 mm x 250 mm, 1.9 μm). High pH separation was performed using a non-linear gradient starting from 4% B to 53% B in 40 min (B: 80% ACN, pH 10.0, adjusted with ammonium hydroxide), 53% B to 68% B in 15 min, 68% B to 95% B in 5 min and maintained here for 3 min. The column was re-equilibrated at the initial condition for 17 min. The column flow rate was maintained at 2 μL/min, and the column temperature was maintained at 30 °C. The EasyPept Frac nano automatic fraction collection system was used to collect 8 fractions, and each fraction was dried in a vacuum concentrator for the next step.

##### DIA Data Collection

The UltiMate 3000 (Thermo Fisher Scientific, MA, USA) liquid chromatography system was connected to the timsTOF Pro 2, an ion-mobility spectrometry quadrupole time of flight mass spectrometer (Bruker Daltonik, Bremen, Germany). Samples were reconstituted in 0.1% FA and 200 ng peptide was separated by AUR3-15075C18 column (15 cm length, 75 μm i.d, 1.7 μm particle size, 120 A pore size) with a 60 min gradient starting at 4% buffer B (80% ACN with 0.1% FA) followed by a stepwise increase to 28% in 25 min, 44% in 10 min, 90% in 10min and stayed there for 7 min, then equilibrates at 4% for 8 min. The column flow rate was maintained at 400 nL/min with a column temperature of 50 °C. DIA data was acquired in the diaPASEF mode. We defined 22 × 40 Th precursor isolation windows from m/z 349 to 1229. To adapt the MS1 cycle time, we set the repetitions to variable steps (2-5) in the 13-scan diaPASEF scheme in our experiment. During PASEF MSMS scanning, the collision energy was ramped linearly as a function of the mobility from 59 eV at 1/K0 = 1.6 Vs/cm^2^ to 20 eV at 1/K0 = 0.6 Vs/cm^2^.

##### DIA Database Search

Raw data were processed and analysed by Spectronaut 18 (Biognosys AG, Switzerland) with default settings. The database used was *Mus musculus* (version2022, 21992 entries), downloaded from Uniprot. Carbamidomethyl on Cysteine was specified as the fixed modification, while oxidation on Methionine was specified as the variable modification. The retention time prediction type was set to ‘dynamic iRT’. Data extraction was determined by Spectronaut based on extensive mass calibration. Spectronaut will determine the ideal extraction window dynamically depending on iRT calibration and gradient stability. Qvalue (FDR) cutoff on precursor level was 1%, and protein level was 1%. Decoy generation was set to ‘mutated’ which is similar to scrambled but will only apply a random number of AA position swamps (min = 2, max = length/2). The normalization strategy was set to ‘local normalization.’ Peptides that passed the 1% Qvalue cut off were used to calculate the major group quantities with the MaxLFQ method.

##### Differential expression analysis

For expression analysis, we used the limma package (v3.54.2) to compare protein expression levels, identifying differentially expressed proteins with an adjusted p-value < 0.05. Following the differential expression analysis, we performed Gene Set Enrichment Analysis (GSEA) using annotated gene sets from the Molecular Signatures Database (MSigDB) to determine whether a predefined set of genes shows statistically significant, concordant differences between two biological states. FGSEA with the fgsea package (v1.28.0) was performed. We then visualized the pathway analysis results through enrichment plots, with an adjusted p-value < 0.05 applied throughout. All analyses were performed using R Statistical Software (v4.2.2; R Core Team 2022).

#### Inductively coupled plasma mass spectrometry (ICP-MS)

##### Sample preparation

All solutions were prepared using ultrapure reagents. Ultrapure water (≥18.2 MΩ cm) from a water purification system (Hydro, Clifton, NJ, USA) was used for reagents and standard solutions. Ultrapure nitric acid (HNO_3_, 65-67%, Optima®) from Fisher Scientific was employed for sample digestion and standard solutions.

Mouse tissues (eWAT, sWAT, and BAT) were prepared for total quantitative analysis of copper (Cu), zinc (Zn), iron (Fe), and selenium (Se) using a microwave-assisted acid digestion method. Samples were defrosted, and wet tissues were weighed with a precision of ±0.01 mg into acid-cleaned 7 mL PFA vials. Subsequently, 0.5 mL of HNO_3_ was added to each vial. For quality control and method assessment, bovine liver (NIST 1577C) from the National Institute of Standards and Technology was used as a reference material. Blank samples consisting only of 0.5 mL of HNO_3_ were also processed (n = 46). The vials were capped and submerged in larger EasyPrep microwave vessels, and the digestion was performed using the MARS 6 Digestion Microwave System (CEM Corp., Matthews, USA). The temperature was ramped up in four stages to 180°C and held for 20 minutes. After cooling, the digested samples were transferred to 15 mL metal-free centrifuge tubes. Then, 50 µL of a gallium (Ga) and yttrium (Y) internal standard mix (500 µg/L stock) was added, and the samples were diluted with ultrapure water to a final volume of 5 mL.

Mouse serum samples were prepared by mixing 0.1 mL of serum with 50 µL of a Ga and Y internal standard solution (500 µg/L stock) in 15 mL metal-free tubes (Labcon, Petaluma, USA). The resulting mixture was then diluted to a final volume of 5 mL with a diluent of 1% HNO_3_, 0.02% Triton X-100, and 500 µg/L gold. For quality control and method validation, certified reference materials (CRMs) QM-S-Q2207 and QM-S-Q2208 from the Quebec Multi-element External Quality Assessment Scheme from the Institut National de Santé Publique (Centre de Toxicologie du Québec, Canada) were used. Method blanks were prepared in the same manner as the serum samples but without the addition of sera.

For the metal analysis of the MSRA immunoprecipitates, 0.1 mL of solution was transferred to 15 mL metal-free tubes and diluted to a final volume of 5 mL with a diluent of 10% HNO_3_ and 500 µg/L gold.

##### ICP-MS measurement

An Agilent 8900 inductively coupled plasma mass spectrometer (ICP-MS) equipped with an Agilent SPS 4 autosampler system was used for the analysis. The standard ICPMS/MS setup employed for all samples included a MicroMist glass nebulizer, a glass double pass spray chamber, platinum/copper sampler and skimmer cones, and a quartz plasma torch with an inner diameter of 2.5 mm. The operating parameters for the ICPMS/MS were as follows: the radiofrequency (RF) power was set to 1,550 W, the plasma gas flow rate was 15.0 L/min, and the auxiliary gas flow rate was 0.9 L/min. The spray chamber temperature was maintained at 2 °C.

External eight-point calibration was performed using matrix-matched solutions. For tissue and MSRA immunoprecipitate samples, the calibration matrix consisted of 10% vol. HNO_3_, 500 µg/L gold, and 5 µg/L Ga and Y internal standard mix. For serum samples, the calibration matrix consisted of 1% vol. HNO_3_, 500 µg/L gold, and 5 µg/L internal standard. Cu, Fe, Se, and Zn were measured in various gas modes. Cu was analyzed in helium mode on mass 63, while Zn was measured on mass 66 in ammonia mode. Fe was measured in oxygen mode, and Se was measured in oxygen mode by a mass shift from 80 to 96. The stabilization time between each gas mode was 10 seconds. The integration times for all masses were set to 0.4 seconds, except for Fe, which had an integration time of 0.1 seconds.

The limit of detection (LOD) was calculated using the formula: LOD = 3.33 × standard deviation from blank measurements. The LODs for microwave-digested tissues were as follows: 3.9 µg/L for Fe, 19 µg/L for Zn, and 0.01 µg/L for Se. For the serum samples, the LODs were 0.2 µg/L for Fe, 0.05 µg/L for Cu, 0.3 µg/L for Zn, and 0.02 µg/L for Se.

The accuracy for CRM NIST 1577c was 96% for Fe, 81% for Cu, 92% for Zn, and 95% for Se, as percentages of certified values. The accuracy for the serum CRMs (QM-S-Q2207 and QM-S-Q2208) was 86-89% for Cu, 102-105% for Zn, and 106-108% for Se, respectively. Fe was not certified for the serum CRMs.

#### Global transcriptome analysis

Total RNA (RIN value > 7) was extracted from adipose tissues, and poly-A pull-down was performed to enrich mRNAs. Library construction was performed using Illumina TruSeq chemistry, with the final PCR step being modified using the KAPA HiFi HotStart Ready Mix. Libraries were sequenced using Element AVITI at the Columbia Genome Center. Samples were multiplexed in each lane, yielding a targeted number of paired-end 75bp reads for each sample. Bases2fastq version 1.4.0.833301531 was used for converting BCL to fastq format, coupled with adaptor trimming. Pseudoalignment to a kallisto index was created from transcriptomes (Mouse:GRCm38.p6) using kallisto (0.44.0). Differentially expressed genes were analyzed using DESeq2 R packages designed to test differential expression between two experimental groups from RNA-seq counts data. Following the differential expression analysis, we performed Gene Set Enrichment Analysis (GSEA) using annotated gene sets from the Molecular Signatures Database (MSigDB) to determine whether a predefined set of genes shows statistically significant, concordant differences between two biological states. For RNA-seq analysis, GSEA with the clusterProfiler package (v4.10.0) was used. We then visualized the pathway analysis results through enrichment plots and dot plots. An adjusted p-value < 0.05 was applied throughout. All analyses were performed using R Statistical Software (v4.2.2; R Core Team 2022).

#### Quantitative RT-PCR

RNA was extracted from snap-frozen adipose tissues or adipocyte cultures using TRIzol reagent (Invitrogen Cat# 15596018). cDNA was synthesized using 1 μg of total RNA and TaqMan Reverse Transcription Reagents (Applied Biosystems Cat# N8080234). All targets were amplified (40 cycles) using gene-specific Taqman probes on a QuantStudio5 Real time-PCR instrument (Applied Biosystems). Relative gene expression quantification was performed using the DDCT method with the QuantStudio Real-Time PCR software v1.1 (Applied Biosystems). Expression levels were normalized to *Hprt*.

##### Taqman probes used

*Hprt* Mm00446968_m1

*MsrA* Mm00452783_m1

*Ucp1* Mm01244861_m1

#### Protein extraction

##### Adipose tissues

Protein extraction from fresh or snap-frozen adipose tissue was performed based on an established protocol (An and Scherer, 2020). In brief, tissues were placed in 500 µL RIPA buffer with protease inhibitor in the absence of Triton X-100 and homogenized until clear. Homogenates were centrifuged at 6,000 x *g* for 15 minutes at 4 °C. The white lipid layer above the aqueous layer was removed, and the pellet was resuspended with Triton X-100 added to a final concentration of 1% (v/v), with 1 tablet of PhosSTOP (Roche Cat# 4906837001) and 1 tablet of cOmplete™, Mini, EDTA-free Protease Inhibitor Cocktail (Roche Cat# 11836170001) per 10 mL buffer, followed by incubation on ice for 60 minutes and clarification at 12 000 x *g* for 15 minutes at 4 °C.

##### Adipocyte culture

Adipocytes were harvested by cell lifters using cold PBS on ice. Protein lysates were prepared using 0.1% SDS lysis buffer in 50 mM Tris pH 8, 0.5% Deoxycholate, 150 mM NaCl, 2 mM EDTA, 1% NP40, with 1 tablet of PhosSTOP (Roche Cat# 4906837001) and 1 tablet of cOmplete™, Mini, EDTA-free Protease Inhibitor Cocktail (Roche Cat# 11836170001) per 10 mL buffer, followed by incubation on ice for 15 minutes and clarification at 12 000 x *g* for 15 minutes at 4 °C.

#### SDS PAGE analysis

Lysates from adipose tissue or cell culture were separated on 4-12% Bis-Tris NuPAGE gels (Invitrogen Cat# NP0335BOX), 12% Bis-Tris SurePAGE Gel (Genscript Cat#M00669; M00667), or 3-8% Tris-Acetate NuPAGE gels (Invitrogen Cat# EA0378BOX), transferred onto nitrocellulose membrane (Amersham Protran 0.45 μm Cat# 10600002) and incubated with the indicated antibodies for immunoblotting. The following primary antibodies were used for immunohistochemical staining at 1:1000 dilution unless stated otherwise: 4-HNE (RRID: AB_722490), ACTIN (RRID: AB_10998774), CS (RRID: AB_2665545), FABP4 (RRID: AB_3082382), FTH1 (RRID: AB_11217441), HSL (RRID: AB_2798800), HSP90 (RRID: AB_2120924), MSRA (1:250, RRID: AB_1853826), MSRB2 (RRID: AB_2813685), MSRB3 (RRID: AB_2720052), Myc-tag (RRID: AB_331783), pHSL-Ser660 (RRID: AB_2893315), PKA (RRID: AB_1523259), PLIN1 (RRID: AB_10829911), pp38-Thr180/Tyr182 (RRID: AB_2139682), pPKA substrates (RRID: AB_331817), TFR1 (RRID: AB_2904534), TH (RRID: AB_297840), TUBULIN (RRID: AB_2288042), UCP1 (1:250, RRID: AB_2241462), VINCULIN (RRID: AB_11458106), Peroxidase-AffiniPure Donkey Anti-Rabbit IgG (RRID: AB_10015282), Peroxidase-AffiniPure Donkey Anti-Mouse IgG (RRID: AB_2340770).

#### Native PAGE analysis

Adipocytes were lysed using 50 mM HEPES pH 7.5, 140 mM NaCl, 1% Triton X-100, and 1% Deoxycholate with 1 tablet of PhosSTOP (Roche Cat# 4906837001) and 1 tablet of cOmplete™, Mini, EDTA-free Protease Inhibitor Cocktail (Roche Cat# 11836170001) per 10 mL buffer. The lysate was mixed with 2X native sample buffer, including 62.5 mM Tris-HCl pH 6.8, 40% glycerol, and 0.01% bromophenol blue, and separated on 16% Novex^TM^ Tricine gels (Invitrogen Cat# EC66952BOX), transferred onto nitrocellulose membrane and incubated with anti-Myc-tag antibody at 1:1000 dilution (RRID: AB_331783).

#### Immunoprecipitations

Cells were lysed with HEPES/IP lysis buffer (550 mM HEPES pH 7.5, 140 mM NaCl, 1% Triton X-100, and 1% Deoxycholate) including 1 tablet of PhosSTOP (Roche Cat# 4906837001) and 1 tablet of cOmplete™, Mini, EDTA-free Protease Inhibitor Cocktail (Roche Cat# 11836170001) per 10 mL buffer on ice and cleared by centrifugation at 16, 000 g for 10 min. Anti-c-Myc-tag magnetic beads (Thermo Fisher Scientific, 88842) were washed three times with HEPES/IP lysis buffer and then incubated overnight with clarified lysates at 4 °C. Beads were washed four times with HEPES/IP lysis buffer and eluted with glycine-HCl buffer (0.1 M, pH 2.2), followed by immediate neutralization by adding Tris buffer (1 M, pH 8.0).

#### *In silico* modeling of oxidized M118 and M120 residues in PKA

All modeling exercises were performed using Maestro (Schro dinger Suite). M118 and M120 residues were individually or collectively modified to their respective oxidized form in the crystal structure of *holo* PKA (PDB code: 3FJQ). This was followed by a preparation workflow with a default setting for global simulation, PROPKA, and heteroatom states at pH 7.4. The resulting hydrogenated models were then minimized using the solvation model VSGB and force file OPLS_2005 as implemented in Maestro.

#### Recombinant protein synthesis

The genes encoding mouse *MsrA* and m*MsrA^AxxA^* were cloned into the pET30a(+) expression vector using NdeI (CATATG) and HindIII (AAGCTT) cloning sites in frame with an N-terminus 6x His-tag. For human recombinant protein, the gene encoding *MSRA* was cloned into the pET30a(+) expression vector with an *N*-terminus 6x His-tag followed by the TEV protease recognition sequence (ENLYFQG). The plasmids were transformed to BL21 StarTM (DE3) chemically competent E. *coli* by heat shock. Transformed cells were grown in LB for 16 h at 15 °C. Cells were lysed, and the expressed proteins were purified through a Ni-NTA HisTrap column. To remove the His-tag, TEV protease was added at a protease-to-substrate ratio of 1:50 (w/w) into His-tagged MSRA in 150 mM NaCl, 0.5 mM TECP, and 20 mM Tris-HCl pH 7.5. The reaction mixture was incubated at 4°C overnight to ensure complete cleavage of the tag. The cleavage reaction mixture was then applied to a second nickel-affinity column to separate the cleaved tag and TEV protease from the untagged target protein MSRA. The flow-through containing the untagged MSRA was collected, followed by concentration and purification by size-exclusion chromatography (SEC) using the Superdex 200 increase column (10/300) (Cytiva, USA).

#### Size exclusion chromatography

All size-exclusion chromatography (SEC) experiments to identify monomeric and dimeric forms of MSRA were performed using a Superdex 200 Increase 5/150 GL (Cytiva, USA) column attached to an Akta Pure (Cytiva, USA), a fast protein liquid chromatography (FPLC) apparatus in a cold room (4 °C). The SEC flow rate was set to 0.1 mL/min using a buffer consisting of 150 mM NaCl, 0.5 mM TCEP, and 20 mM Tris-HCl, pH 7.5. Every SEC run was monitored by the Akta Pure UV detector at the wavelength 280 nm. The eluted samples were collected in 100 μL fractions for further biochemical and biophysical analyses.

#### Methionine sulfoxide reductase activity assay

For MSRA activity analysis, a reaction mixture (50 µL) containing 20 mM *L*-methionine, 20 mM *L*-methionine sulfoxide, 30 mM KCl, 10 mM MgCl_2_, and 20 mM DTT was prepared. SEC-purified recombinant MSRA monomer or dimer was added, and the mixture was incubated at 37 °C for 30 min. After adding 300 µL ddH_2_O, MSRA protein was removed using Amicon® ultra 0.5 mL 10k centrifugal filters with 10,000 x *g*. Reverse-phase HPLC determined the *L*-methionine and *L*-methionine sulfoxide abundance in the filtrate with UV detection at 214 nm (Agilent 1260 Infinity II). Water (solvent A) and acetonitrile (solvent B) containing 0.1% TFA were used as solvents to make a gradient in HPLC. HPLC separation was performed at 1 mL min^−1^ flow rate using the following linear gradient: 0–2 minutes: 0% B, 2–32 minutes: 0–70% B, 32–33 minutes: 70–100% B, 33–35 minutes: 100% B, 35–36 minutes: 100–0%. Retention times for *L*-methionine and *L*-methionine sulfoxide were 3.8 min and 2.1 min, respectively.

#### Oxaziridine Synthesis and NMR validation

Oxaziridine-alkyne (Ox-alkyne) (He et al., 2022; Lin et al., 2017) was synthesized based on published procedures. ^1^H-NMR and high-resolution mass spectrometry validated synthesized oxaziridine-alkyne. ^1^H NMR (400 MHz, MeOD): δ (ppm) = 7.55–7.40 (m, 5H), 5.09 (s, 1H), 4.22 (t, 2H, J = 1.6 Hz), 3.61 (s, 2H), 2.89 (1H), 1.38 (s, 6H). HR-MS: calcd. for C_15_H_18_O_3_N_2_Na [M + Na]^+^, 297.1215; measured: 297.1208.

#### Oxaziridine-alkyne/Iodoacetamide-alkyne labeling

Oxaziridine-alkyne (100 μM) and iodoacetamide-alkyne (100 μM) labeling were conducted on freshly prepared cell lysates or mouse adipose tissue lysates for 1 hour at 25 °C. For endogenous protein, copper-catalysed azide-alkyne cycloadditions (CuAAC) were performed using 200 μM azide-(PEG)_3_-biotin probe (Sigma-Aldrich Cat# 762024), followed by CHCl_3_/MeOH precipitation. The precipitated protein samples were solubilized in 2% SDS in PBS and then diluted to 0.1% SDS for subsequent procedures. The solutions were added to streptavidin Sepharose high-performance beads (Cytiva Cat# 17511301) and incubated at 4 °C overnight with rotation. After washing with PBS three times, proteins were eluted from streptavidin agarose gel by adding 1X LDS sample buffer (Invitrogen Cat# NP0007), boiled at 95 °C for 5 min, and blotted for proteins of interest.

#### Quantifications and Statistical Analysis

Biochemical experiments *in vitro* were repeated at least three times, and the repeat number was increased according to effect size or sample variation. We estimated the sample size considering the variation and mean of the samples. No statistical method was used to predetermine the sample size. No animals or samples were excluded from any analysis. Animals were randomly assigned groups for *in vivo* studies; no formal randomization method was applied when assigning animals for treatment. All western blotting experiments with quantification were performed a minimum of three times with independent biological samples and analyzed by ImageJ 1.52q. Investigators were blinded to group allocation during data analysis. Statistical analyses were performed using GraphPad Prism 8. All tests and p values are provided in the corresponding Figs. or Fig. legends.

