## Supplementary Figure Legends for "An Iron-regulated Signalling Pathway Controls Adipose Browning and Cancer Cachexia"

**Supplementary Figure 1.**

**a**, Normalized fat mass of pancreatic tumor-bearing mice (mPDA) compared to age-matched non-tumor controls (mNT). F, female; M, male.

**b**, qRT-PCR of *Ucp1* mRNA expression in epididymal eWAT of pancreatic tumor-bearing mice (mPDA) compared to age-matched non-tumor controls (mNT).

**c**, Immunohistochemical analysis of UCP1 expression in BAT and WAT pancreatic tumor-bearing mice (mPDA) and age-matched non-tumor controls (mNT).

**d-f**, Locomotor activity (**d**), food intake (**e**), and water intake (**f**) of pancreatic tumor-bearing mice (mPDA) compared to age-matched non-tumor controls (mNT). Welch’s *t*-test was performed, given a significant variation in the variance of locomotive activity, food, and water intakes of tumor-bearing mice compared to non-tumor controls (F-test, p < 0.01 for each parameter).

**g**, Immunohistochemical analysis of tyrosine hydroxylase expression in BAT and sWAT of pancreatic tumor-bearing mice (mPDA) compared to age-matched non-tumor controls (mNT).

**h**, Gene Set Enrichment Analysis of the global proteome of sWAT from female pancreatic tumor-bearing mice (mPDA) compared to age-matched non-tumor controls (mNT).

**i,** Gene Set Enrichment Analysis of the global proteome of sWAT from non-tumor-bearing male mice treated with 5 mg/kg β3-adrenergic agonist CL-316243 (CL) compared to vehicle (mNT) for 6 days.

**j**, Immunoblot analysis of 4-hydroxynonenal (4-HNE) modified proteins in sWAT of pancreatic tumor-bearing mice (mPDA) compared to age-matched non-tumor controls (mNT). Right, quantification of 4-HNE levels normalized to protein input.

**k**, Immunoblot analysis of protein expression in eWAT of pancreatic tumor-bearing mice (mPDA) compared to age-matched non-tumor controls (mNT)

**l**, Immunoblot analysis of protein expression in BAT, eWAT, and sWAT from non-tumor-bearing mice. WT, *MsrA^+/+^*; KO, *MsrA^–/–^*.

**m**, Immunoblot analysis of protein expression in sWAT from male non-tumor-bearing mice treated with 5 mg/kg β3-adrenergic agonist CL-316243 (CL) or vehicle for 6 days.

**n**, Adipocyte diameter of white adipose tissues from non-tumor patients who underwent laparoscopic cholecystectomy (hNT) or were diagnosed with cachexic PDA (hPDA). 100 adipocytes were measured per specimen. Means ± SEMs.

**o**, Immunohistochemical analysis of UCP1, TNAP and COX4 expression in the visceral WAT from non-tumor patients who underwent laparoscopic cholecystectomy (hNT) or were diagnosed with cachexic PDA (hPDA).

**p**, Immunoblot analysis of WAT from pancreatic cancer patients with (hPDA^CAC^, body mass loss >5%) or without (hPDA^nonCAC^, no or < 5% body mass loss) cachexia.

Error bars in this Fig. are means ± SDs unless stated otherwise. Student’s *t*-test was performed unless stated otherwise. N.S., not significant.

**Supplementary Figure 2.**

**a**, Immunoblot analysis of protein expression in eWAT and sWAT from pancreatic tumor-bearing mice (mPDA) compared to age-matched non-tumor controls (mNT).

**b**, Immunoblot analysis of protein expression in WAT from non-tumor patients who underwent laparoscopic cholecystectomy (hNT) or were diagnosed with cachexic PDA (hPDA).

**c**, Total iron (Fe) and zinc (Zn) levels in the adipose tissues of pancreatic tumor-bearing mice (mPDA) compared to age-matched non-tumor controls (mNT).

**d**, Total iron (Fe) and zinc (Zn) levels in the serum of pancreatic tumor-bearing mice (mPDA) compared to age-matched non-tumor controls (mNT).

**e**, Representative Hematoxylin and Eosin (H&E) staining of brown (BAT) and subcutaneous white (sWAT) adipose tissues from mice treated with vehicle, 5 mg/kg β3-adrenergic agonist (CL-316243), 50 mg/kg deferiprone (DFP), or in combination (CL+DFP) for 6 days.

**f**,**g**, qRT-PCR (**f**) and immunoblot (**g**) analysis of white adipocyte cultures expressing control sgRNA (sg*Rosa*) or sgRNA against the ferritin heavy chain (sg*Fth1*). Right, quantification of MSRA protein expression relative to loading control VINCULIN (VIN).

**h**, Inductively coupled plasma mass spectrometric analysis of total iron and zinc content in MSRA immunoprecipitants in the presence or absence of exogenous ferric ammonium citrate (FAC).

Error bars in this Fig. are means ± SDs. Student’s *t*-test was performed. N.S., not significant.

**Supplementary Figure 3.**

**a**, Metal Ion-Binding site prediction of MSRA (left) and Ribonucleotide Reductase (right).

**b**, Native PAGE analysis and quantification of MSRA expression in white adipocyte cultures in the presence of exogenous ferric ammonium citrate (FAC) or iron chelator deferiprone (DFP).

**c**, Size exclusion chromatogram of recombinant wildtype (WT) and iron binding domain mutant (AXXA) of mouse MSRA in the presence or absence of 5 molar equivalents of ferric ammonium citrate (FAC). Inset, Coomassie analysis of the recombinant proteins used in the analysis.

**d-g**, Activity assay of increasing concentrations of gel filtration purified mouse MSRA dimer (**d**) and monomer (**e**), quantifications of Met/Met-O ratio (**f**), and comparison to unfractionated MSRA-WT (**g**).

**Supplementary Figure 4.**

**a**, Immunoblot analysis of WAT from pancreatic cancer patients with (hPDA^CAC^, body mass loss >5%) or without (hPDA^nonCAC^, no or < 5% body mass loss) cachexia. The level of PKA substrate phosphorylation is quantified relative to the total PKA expression.

**b**, Schematic of oxazaridine labelling to measure proteinaceous methionine reactivity.

**c**, Structural overlay of *holo* PKA (cyan, PDB code 3FJQ) with ADP-bound PKA (olive, PDB code 1JBP). ATP (black for carbon atoms), ADP (dark pink for all atoms), and P-loop (magenta) are labeled.

**d**, Structural overlay of *holo* WT PKA (cyan, PDB code 3FJQ) and molecular dynamic (MD) simulated model (Schrodinger Suite) of two oxidized methionine residues of PKA at position 118 (SME118) and 120 (SME120). The cofactor ATP is shown as stick models for WT PKA (in black) and the MD simulated model when M118 and M120 are oxidized (in brown).

**e**, Phenix refined structures of oxidized M118 and M120 using experimental data obtained from the *holo* PKA (PDB code 3FJQ). Steric clashes on the surrounding residues between oxidized M118 (SME118) or M120 (SME120) or together are shown with the red cloud.

**f**, Representative Hematoxylin and Eosin (H&E) staining of brown (BAT) and white (WAT) adipose tissues from non-tumor-bearing mice.

**g**, Immunoblot analysis of protein expression from BAT and WAT of non-tumor-bearing mice.

**h**, Brightfield images of preadipocyte cultures as they differentiate into white adipocytes.

**i**, Immunoblot analysis of PKA protein expression in white adipocyte cultures upon treatment with the pan-β-adrenergic receptor agonist (isoproterenol; ISO) or the PKA inhibitor (H89) at the indicated concentrations and times.

**j**, Immunoblot analysis of activating phosphorylation of hormone lipase (pHSL) in white adipocyte cultures upon treatment with the pan-β-adrenergic receptor agonist (isoproterenol; ISO) or the PKA inhibitor (H89) at the indicated concentrations and times.

WT, *MsrA^+/+^*; KO, *MsrA^–/–^*. Error bars in this Fig. are means ± SDs. Student’s *t*-test was performed. N.S., not significant.

**Supplementary Figure 5.**

**a**, Body mass of tumor-bearing mice compared to age-matched non-tumor controls. KPC, *Kras^G12D^;p53^R172H^;PdxCre;MsrA^+/+^*; KPCM, *Kras^G12D^;p53^R172H^;PdxCre;MsrA^–/–^;* WT, non-tumor *MsrA^+/+^*; KO, non-tumor *MsrA^–/–^.*

**b**, Percentage change of body mass over 4 weeks in pancreatic-tumor-bearing mice upon the detection of palpable tumors.

**c**, Immunoblot analysis of MSRA expression in tibial skeletal muscles of age-matched tumoring-bearing mice compared to age-matched non-tumor controls (mNT).

**d**, Lean mass of tumor-bearing mice compared to age-matched non-tumor controls.

**e**, Skeletal muscle cell diameter of tumor-bearing mice compared to age-matched non-tumor controls (mNT). 100 myocytes were measured per specimen. Means ± SEMs.

**f**, The mass of pancreatic tumors from moribund mice.

**g**, Representative Hematoxylin and Eosin (H&E) staining of pancreatic tumor (PDA) and other organs from tumor-bearing mice.

**h**, Light and dark cycle food intake.

**i**, Immunoblot analysis of tyrosine hydroxylase (TH) expression in the sWAT of pancreatic tumor-bearing mice.

**j**, Immunohistochemical analysis of β3-adrenergic receptor (β3-AR) expression in the sWAT of pancreatic tumor-bearing mice.

Error bars in this Fig. are means ± SDs unless indicated otherwise. Student’s *t*-test was performed. N.S., not significant.

**Supplementary Figure 6.**

**a**, Relative body mass of male mice after orthotopic transplantation of pancreatic cancer cells (mPDA) compared to sex and age-matched non-tumor controls (mNT). Each line represents one animal.

**b**, Correlation of survival and the rate of body mass change over the course of tumor development in male mice orthotopically transplanted with pancreatic cancer cells.

**c**, Pancreatic tumor, fat, and lean mass normalized to body mass between 3 to 6 weeks after orthotopic implantation of pancreatic cancer cells.

**d**, Gene Set Enrichment Analysis of the sWAT transcriptome of male mice after orthotopic transplantation of pancreatic cancer cells compared to age-matched non-tumor male controls.

**e,** Top 20 upregulated genes in the sWAT of male mice after orthotopic transplantation of pancreatic cancer cells compared to age-matched non-tumor male controls.

**f**, Skeletal muscle cell diameter of male mice 5 weeks after orthotopic transplantation of pancreatic cancer cells, compared to age-matched non-tumor male controls (mNT). WT-host, *MsrA^+/+^*; KO-host, *MsrA^–/–^*. 100 myocytes were measured per specimen. Means ± SEMs.

**g**,**h**, Tumor burden (**g**) and metastatic incidence (**h**) of male mice 5 weeks after orthotopic transplantation of pancreatic cancer cells.

**i**, Immunoblot analysis of tyrosine hydroxylase (TH) expression in the sWAT from male mice 5 weeks after orthotopic transplantation of pancreatic cancer cells.

**j**, Inductively coupled plasma mass spectrometry (ICP-MS) to measure total iron content in adipose tissues from male mice 5 weeks after orthotopic transplantation of pancreatic cancer cells.

**k**, Immunoblot analysis of MSRB2 and MSRB3 expression in the sWAT from male mice 5 weeks after orthotopic transplantation of pancreatic cancer cells, compared to age-matched non-tumor controls (mNT). Loadin control, Tubulin, is shared with Figure 4h.

**l**, Immunoblot analysis of MSRB2 expression in the sWAT from PDA-bearing mice compared to non-tumor controls (mNT). KPC, *Kras^G12D^;p53^R172H^;PdxCre;MsrA^+/+^*; KPCM, *Kras^G12D^;p53^R172H^;PdxCre;MsrA^–/–^*.

Error bars in this Fig. are means ± SDs unless indicated otherwise. Student’s *t*-test was performed. N.S., not significant.

**Supplementary Figure 7.**

**a**, Tumor burden of moribund male mice after orthotopic transplantation of pancreatic cancer cells. WT-host, *MsrA^+/+^* (n = 8); KO-host, *MsrA^–/–^* (n = 7).

**b**, Metastasis incidence of moribund male mice after orthotopic transplantation of pancreatic cancer cells. WT-host, *MsrA^+/+^* (n = 8); KO-host, *MsrA^–/–^* (n = 7).

Error bars in this Fig. are means ± SDs. Student’s *t*-test was performed. N.S., not significant.

**SUPPLEMENTARY TABLE LEGENDS**

**Supplementary Table 1.** Liquid chromatography-mass spectrometry in Data Independent Acquisition (DIA) mode analysis of the proteome of subcutaneous white adipose tissues from mice.

**Supplementary Table 2.** Human specimen characteristics. This Excel file contains the biometrics of each deidentified human patient from whose visceral white adipose tissues were used in this study. hNT, patients who underwent laparoscopic cholecystectomy, and serve as non-tumor controls. hPDA^CAC^, patients diagnosed with PDA exhibit unintentional body mass loss equal to or greater than 5% in 6 to 12 months. hPDA^nonCAC^, patients diagnosed with PDA and exhibit no or less than 5% body mass loss in 6 to 12 months.

**Supplementary Table 3.** RNA-seq transcriptomic analysis of subcutaneous white adipose tissues from mice.
